# Neurophysiological signatures of duration and rhythm prediction across sensory modalities

**DOI:** 10.1101/183954

**Authors:** Acer Y.-C. Chang, Anil K. Seth, Warrick Roseboom

**Affiliations:** Department of Informatics, University of Sussex, Brighton BN1 9QJ, United Kingdom; Sackler Centre for Consciousness Science, University of Sussex, Brighton BN1 9QJ, United Kingdom; Department of Neuroinformatics, Araya, Tokyo, Japan

## Abstract

Effective behaviour and cognition requires the ability to make predictions about the temporal properties of events, such as duration. It is well known that violations of temporal structure within sequences of stimuli lead to neurophysiological effects known as the (temporal) mismatch negativity (TMMN). However, previous studies investigating this phenomenon have typically presented successive stimulus intervals (i.e., durations) within a rhythmic structure, conflating the contributions of rhythmic temporal processing with those specific to duration. In a novel behavioural paradigm which extends the classic temporal oddball design, we examined the neurophysiological correlates of prediction violation under both rhythmically (isochronous) and arrhythmically (anisochronous) presented durations, in visual and auditory modalities. Using event-related potential (ERP), multivariate pattern analysis (MVPA), and temporal generalisation analysis (TGA) analyses, we found evidence for common, and distinct neurophysiological responses related to duration predictions and their violation, across isochronous and anisochronous conditions. Further, using TGA we could directly compare processes underlying duration prediction violation across different modalities, despite differences in processing latency of audition and vision. We discovered a common set of neurophysiological responses that are elicited whenever a duration prediction is violated, regardless of presentation modality, indicating the existence of a supramodal duration prediction mechanism. Altogether, our data show that the human brain encodes predictions specifically about duration, in addition to those from rhythmic structure, and that the neural underpinnings of these predictions generalize across modalities. These findings support the idea that time perception is based on similar principles of inference as characterize ‘predictive processing’ theories of perception.

## Introduction

Adaptive behaviour requires the ability to make predictions about when events will occur, which involves the perception of temporal properties such as duration. There is a long history of treating perception in terms of hypothesis-testing or inference (Gregory, 1968; Helmholtz, 1867), with this idea enjoying recent prominence in terms of the predictive processing approach to perception, action, and cognition (Clark, 2012; Friston, 2005; Seth, 2014). Much of the corresponding experimental work has focused on isolating the neural correlates of prediction and/or prediction error. One classic example is based on neurophysiological event-related potential (ERP) component called the mismatch negativity (MMN) (Friston, 2005; Garrido, Kilner, Stephan, & Friston, 2009; Lieder, Stephan, Daunizeau, Garrido, & Friston, 2013). The MMN is a neural response that can be found following experience of infrequent or *unexpected* events (or their absence) in a sequence.

The classical MMN was demonstrated using rhythmic sequences of auditory events wherein most presentations, standard trials (e.g. 80%), are of one pitch (e.g. low pitch) and infrequently (20%) a deviant or oddball pitch (e.g. high pitch). Under these conditions, deviant trials elicit a distinct ERP waveform as compared to standard trials, characterising the MMN (Näätänen, Gaillard, & Mäntysalo, 1978; Näätänen & Michie, 1979; Sams, Paavilainen, Alho, & Näätänen, 1985). Similar findings have been reported when the deviant is temporal rather than featural such that the rhythm of a sequence of events is disrupted by one event occurring earlier or later than expected based on the rhythm – a temporal oddball paradigm *(temporal* MMN; TMMN, Chen, Huang, Luo, Peng, & Liu, 2010; Jacobsen & Schröger, 2003; Näätänen, Paavilainen, & Reinikainen, 1989; Tse & Penney, 2006; see also review Ng & Penney, 2014).

Interpretation of the TMMN in many studies is complicated by the use of fixed inter-trial-intervals (ITI) (Hsu et al., 2010; Jacobsen & Schröger, 2003, 2003; Joutsiniemi et al., 1998; Okazaki, Kanoh, Takaura, Tsukada, & Oka, 2006; Roger, Hasbroucq, Rabat, Vidal, & Burle, 2009; Takegata, Tervaniemi, Alku, Ylinen, & Näätänen, 2008; Tse & Penney, 2006; for non-fixed ITIs see also Chen et al., 2010; Grimm et al., 2006; Grimm, Widmann, & Schröger, 2004). Fixed ITIs create a rhythmic structure, leaving open the possibility that the TMNN is related to violation of rhythmic structure, rather than violation of duration predictions more specifically. Indeed, deviant rhythmic structures have been shown to cause mismatch responses in both ERPs (Ford & Hillyard, 1981; Vuust et al., 2011) and event-related fields (ERF) recorded by magnetoencephalography (Vuust, Ostergaard, Pallesen, Bailey, & Roepstorff, 2009).

To investigate this possibility, we compared mismatch responses in isochronous and anisochronous stimulus sequences, in which the ITI was either fixed, or was sampled from a distribution that removes global rhythmic structure. We also examined whether neural signatures related to duration prediction are modality-general or specific. Some studies have suggested that duration processing might be modality specific (Wearden, Todd, & Jones, 2006). However, accurate temporal predictions may benefit from the integration of temporal information from different sources (Shi, Church, & Meck, 2013). If so, we would expect to find similar neural activation patterns related to duration prediction across auditory and visual domains, despite well-known differences in sensory evoked potentials.

To examine these hypotheses, we recorded EEG while human participants were passively exposed to isochronous (auditory) or anisochronous (auditory and visual) sequences of durations in which duration oddballs were embedded. We used ERP and multivariate pattern analysis (MVPA) to characterize neural response patterns related to duration prediction. We also used temporal generalisation analysis (TGA; J.-R. King & Dehaene, 2014) to investigate whether the neural signatures of prediction violation in a given condition (e.g. anisochronous auditory sequences) could be used to decode prediction violations in other conditions (e.g. isochronous auditory or anisochronous visual sequences) – overcoming differences in sensory processing latency between modalities that previously made this comparison impossible. Altogether, this combination of analyses allowed us to examine the commonalities and differences in neural responses related to duration prediction between rhythmic and arrhythmic conditions, and across sensory modalities.

## Methods

### Participants

16 healthy students with normal or corrected-to-normal vision were recruited from the University of Sussex (8 males, age range 18-31 years). Written informed consent was acquired from all participants prior to the study, which was approved by the University of Sussex ethics committee. Participants received £15 as compensation for their time.

### Procedure and Design

Participants were seated in a dimly lit, electromagnetically shielded room and asked to maintain fixation ~ 50 cm away from a gamma corrected LaCie Electron blue IV 22" CRT Monitor. Stimuli were generated and presented using the Psychophysics toolbox (Brainard, 1997; Kleiner, Brainard, & Pelli, 2007). Auditory stimuli were played through stereo speakers with an intensity level of ~ 65 dB SPL.

To investigate the neural signatures of duration prediction, we extended a temporal oddball paradigm. In each trial, two transient stimuli were presented, the first stimulus (S1) defining the beginning and the second stimulus (S2) the end of the specified duration (i.e., inter-stimulus-interval, ISI). The duration could be either 150ms or 400ms (Figure 1). There were two blocks of trials for each experimental condition. In one block of trials, a duration of 150ms was presented in 200 trials (standard) with a duration of 400ms presented in 50 trials (deviant). In the other block of trials, the standard and deviant duration were switched. The position of the deviants within the sequence was pseudo-randomised but constrained so that the deviant was presented at least once in every five trials.

**Figure 1.**
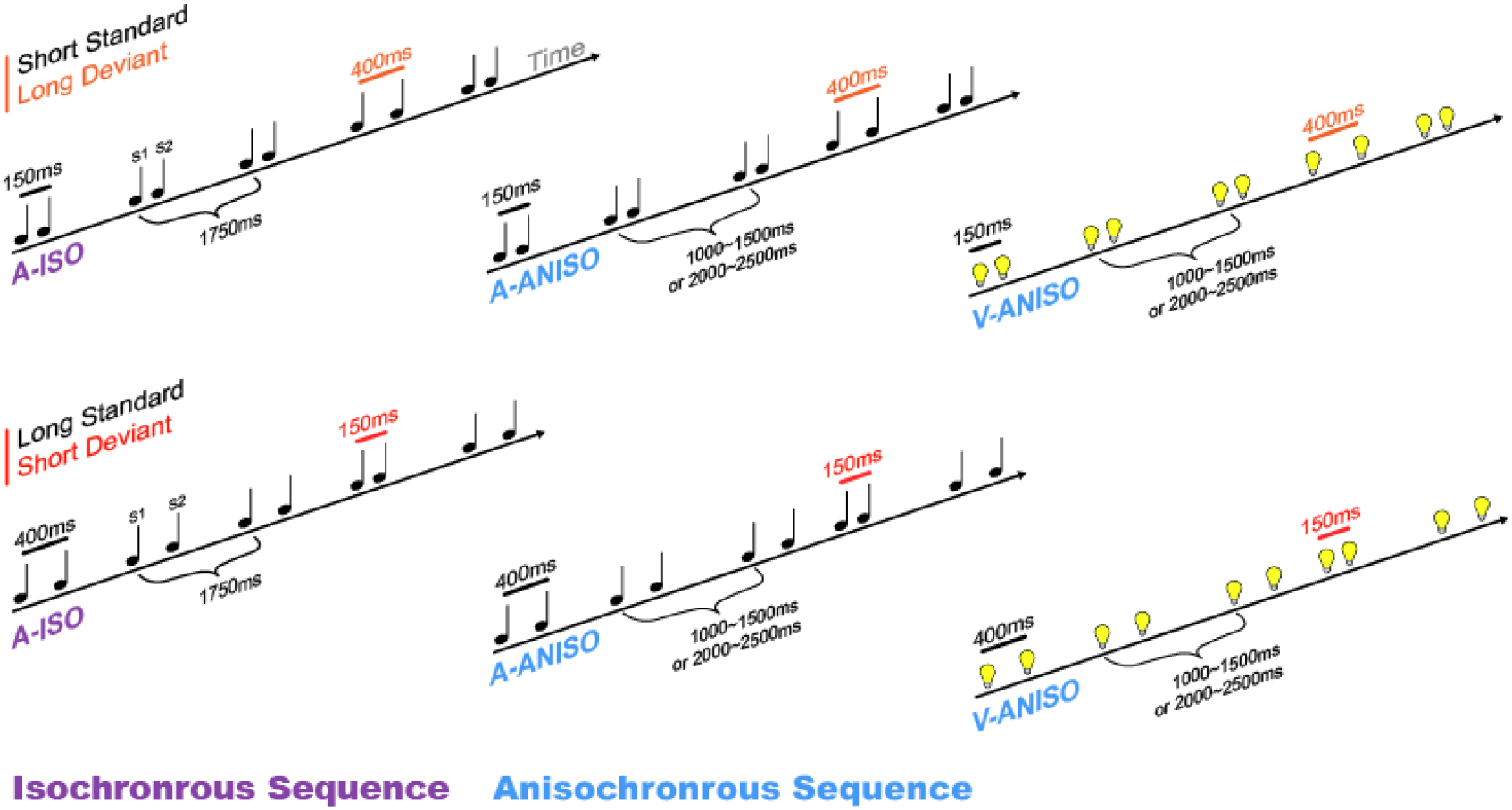
Example stimulus sequences. Presented durations were either short (150ms) or long (400ms). In one experimental block, the standard was short and the deviant long (top row), while in another block this was reversed (bottom row). Isochronous sequences (A-ISO, left column) had a fixed inter-trial-interval (ITI) – the time between onsets of successive durations was always 1750ms. Anisochronous sequences (A-ANISO, the middle column, and V-ANISO, the right column) had a pseudo-random ITI drawn from two uniform distributions two uniform random distributions between 1000-1500ms and 2000-2500ms, and so never included ITIs close to the isochronous ITI of 1750ms. Anisochronous sequences were presented using either auditory (A-ANISO) or visual (V-ANISO) stimulus sequences.

Three experimental conditions were examined in this study: auditory isochronous (A-ISO), auditory anisochronous (A-ANISO), and visual anisochronous (V-ANISO). In the auditory conditions stimuli consisted of 10ms pulses of 1500 Hz pure tones. In the visual condition stimuli were 10ms flashes of luminance-defined Gaussian blobs against a grey background (Michelson contrast of 1). The inter-trial-interval (ITI, the time between the first event in one pair and the next) in the isochronous condition was fixed at 1750ms. In the anisochronous conditions, the ITIs were drawn from two uniform random distributions between 1000-1500ms or 2000-2500ms so that the average presentation rate was the same as the ITIs in the isochronous condition, but never included an ITI equivalent to that from the isochronous condition (1750ms) (Figure 1).

There were no explicit task requirements. Participants were simply instructed to attend to the sequence of events. The order of completion of the six blocks was pseudo-randomised across participants. Each block of 250 trials took approximately 10 minutes to complete and the entire experiment took approximately 1 hour.

### EEG Acquisition

All EEG data were recorded using a 64 channel ANT Neuro amplifier at a sampling rate of 2048 Hz. A 64 channel Waveguard EEG cap (ANT Neuro, Enschede) employing standard Ag/AgCl electrodes placed according to the 10-20 system was used. Horizontal and vertical eye movements were recorded by two independent pairs of electrodes. The impedances of recording electrodes were maintained below 10kΩ. No analog filter was applied during on-line recording.

### EEG Preprocessing

Pre-processing was performed using the EEGLAB toolbox (Delorme & Makeig, 2004) under Matlab (Mathworks, Inc. Natick, MA, USA), and custom Matlab scripts. Continuous data were first down-sampled to 512 Hz. The signals were then band-pass filtered at 0.5-30 Hz and epoched between -140ms and 1400ms relative to onset of the first stimulus in each stimulus pair. Each epoch was baseline corrected and artefacts were automatically identified and removed using the automated Independent Component Analysis (ICA) rejection algorithms ADJUST (Mognon, Jovicich, Bruzzone, & Buiatti, 2011) and MARA (Winkler, Haufe, & Tangermann, 2011). Epochs containing signal values exceeding ±75 or 6 standard deviations from the mean on any single channel were automatically rejected. For the multivariate pattern analyses (MVPA, mentioned below), epochs were further down-sampled to 128 Hz.

### ERP Analysis

For each ERP analysis, epochs were averaged separately within each comparison condition for each participant. Statistical analyses of the ERP data were performed at the group level (16 participants, see Statistical Analyses and significance testing)

### Multivariate Pattern Analysis (MVPA)

#### Decoding EEG activation patterns between standard and deviant evoked responses

To test whether standard and deviant durations elicited distinct topographical patterns in EEG responses, we applied MVPA to EEG topography. As with other applications of MVPA, we reasoned that if the topographical patterns between standard and deviant trials are distinguishable across conditions, a binary classifier should be able to classify topographic patterns recorded from different conditions and reach above chance classification performance. We employed a linear Support Vector Machine (SVM) classifier (Vapnik, 2013) for every classification analysis. In the current study, an unbalanced trial number between standard and deviant trials (close to 4:1; the exact ratio depending on the trials excluded during pre-processing) was expected because of the nature of the oddball paradigm. To avoid biased classification due to training on unbalanced data sets, every MVPA analysis was performed on subsampled data sets (Maimon & Rokach, 2005). A subsampled data set consisted of all deviant trials and randomly subsampled standard trials with equal trials number of deviant trials (mean 45.14 trials).

For each time point and each participant, we performed the MVPA on the data from 64 channels as features in two steps. First, to find the optimized regularization parameter (C: Cost), a subsampled data set was selected and normalised to z-scores (mean subtracted and divided by standard deviation within features, i.e., electrodes) to examine the optimized C. We searched the parameter space from 2^-3 to 2^3 (exponentially spaced) and evaluated the classification performance (classification accuracy) by stratified ten-fold cross-validation (CV). For each fold of the ten-fold CV the subsampled data set was split into training and testing trials with a 9:1 ratio. An SVM classifier was fit on the training trials (training set) and tested on the testing trials (testing set). The classification accuracies across ten folds were than averaged as the CV accuracy. The C that maximised the CV accuracy was selected as the optimal C value.

In the second stage, a new subsampled data set was used. The new subsampled data set contained no overlapping standard trials from the first step. We performed the same CV procedure as the first step except that the C was fixed at the optimal value obtained from the first step. The CV accuracy was then computed as the classification accuracy. We repeated this procedure 50 times with different subsampled data sets in each repetition. The classification accuracies across repetitions were then averaged to obtain a stable and unbiased classification performance across entire data set. All classification analysis were conducted by LIBSVM library (Chang & Lin, 2001) included in e1071 and Caret packages in R software (Kuhn, 2008; Meyer, Dimitriadou, Hornik, Weingessel, & Leisch, 2015).

### Temporal generalization analysis

To examine whether durations presented in isochronous and anisochronous sequences, and auditory or visual modalities, share common underlying neural processes, we employed temporal generalisation analysis (TGA) (adapted from King & Dehaene, 2014). The standard MVPA decoding analysis (above) is adequate to assess whether EEG signals are informative for successfully distinguishing standard and deviant evoked activation patterns at each analysis time point. However, because information processing may evolve differently between conditions, an analysis approach that accommodates potential latency differences between conditions is needed. For this purpose, TGA was used to evaluate common processing between conditions. If a classifier trained on activation patterns at a specific time point can also classify activation patterns at another time point with above chance accuracy, one can infer the existence of similar underlying neural processes at the two time points. Therefore, TGA provides an estimation of common processing across time, and can be additionally used to examine the latency difference of common processes between conditions (J.-R. King & Dehaene, 2014, p. 2).

As with the MVPA analysis, the TGA procedure was repeated 50 times (runs) for each training time point and participant. In each run, a subsampled dataset containing an equal number of standard and deviant trials (the training set) for a given training time point (training time, i.e., t_training_) in one condition was trained. This classifier was then used to test subsampled, balanced data sets in which each data set (testing sets) was the data from a time point (testing time, i.e., t_testing_) in another condition. Therefore, a trained linear SVM model from one condition was used to predict the trial types (standard or deviant) at time points in the data from another condition. For example, one can train a classifier using EEG data at 300ms (training set) in an auditory anisochronous sequence to classify standard and deviant trials. If the trained classifier is able to successfully predict the trial types using data at 350ms (testing set) in auditory anisochronous sequence, this suggests that similar decodable information is embedded in the training and testing sets and can be captured by the classifier even from different time points and conditions. Overall, this procedure yielded 198 generalisation accuracies (1.54s epoch length x 128Hz sampling rate = 198 sampling points). The same procedure was applied to each training time point (198 training time point in total). A generalization matrix with training time x testing time (198 x 198) was obtained to examine the possible shared neural processes between conditions in each run. Finally, the 50 generalisation matrices from the 50 runs were then averaged.

In a generalization matrix, an above chance generalisation accuracy on the diagonal (training time = testing time) suggests shared processes occurring at the same latencies between conditions. If a matrix shows an off-diagonal pattern, the shared processes occurs at different timings in the two conditions (J.-R. King & Dehaene, 2014; J.-R. King, Gramfort, Schurger, Naccache, & Dehaene, 2014).

### Statistical analyses and significance testing

Cluster-based permutation analysis with Monte Carlo randomization (Maris & Oostenveld, 2007) was performed for both ERP and MVPA analyses using custom scripts adapted from FieldTrip (Nichols & Holmes, 2002; Oostenveld, Fries, Maris, & Schoffelen, 2010) and the Mass Univariate Toolbox (Groppe, Urbach, & Kutas, 2011). This method was conducted to reduce the family-wise type I error rate due to multiple comparisons. The algorithm considers an effect is a true positive when a group of adjacent analysis points (neighbours) reach statistical significance together. Two steps were taken in this procedure. First, for every testing point, a statistic across participants was computed (e.g. t-value). A critical value was then used for thresholding testing (e.g. p < .05, one-tail). Second, adjacent data points that exceeded the critical value together were defined as a cluster. For each cluster, the cluster ‘mass’ was computed by summing all statistics from each data point within a given cluster. To construct a null distribution from permutations, the same procedure was repeated 5000 times with shuffled condition labels for each data point and participant. The maximum cluster mass from each permutation was then used to construct the null distribution. Finally, the cluster-level p-value was computed by identifying the rank of cluster mass from real data in the null distribution (Monte Carlo p value).

For ERP analyses, the neighbours were defined as adjacent channels and adjacent sample time points. The adjacent channels were computed by Delaunay triangulation in 2D projection of the sensor position. In the first step, critical values were computed using a two-tailed dependent t statistic with a p < .05 criterion.

For, MVPA analyses for 1D data (time only), the neighbours were defined as adjacent sample time points. For the TGA, the neighbours were defined as adjacent training and testing time points in a 2D space. Because we only considered above chance level classification accuracy, critical values were computed from one-tail, one-sample t statistics again using 50% accuracy with a p < .05 criterion.

For the robust multilinear regression analysis, a one sample t-test was performed to test the Fisher’s z-score for each time point against an average score calculated from the pre-stimulus period.

## Results

### Neural correlates of duration prediction

#### Prediction violation in isochronous auditory sequences

Neural correlates of duration prediction were examined by identifying neural responses to sensory events that occurred before the predicted temporal interval had elapsed. In a long standard duration block of trials, participants were presented with the long standard duration (400ms) and occasional short duration (150ms) deviants. In such deviant trials, the second stimulus (S2) arrived earlier than would be predicted (150ms). By comparing the short duration deviant trials (150ms) with physically identical short duration standard trials (150ms), presented in the counterbalanced block, we can examine neural responses to an unexpected duration with physically identical sensory stimulation.

We first replicated the results of previous temporal oddball studies in which rhythmic information, due to fixed ITIs (isochronous sequences), may also have contributed to temporal prediction of sensory events. The results from ERP comparison between standard and deviant short duration trials in A-ISO sequences showed two pairs (both significant positive and negative parts within similar time windows) of significant clusters (Figure 2, A-ISO). The early paired clusters at 227ms-313ms showed a negative ERP deviation in response to the unexpected short duration, with a central to frontal distribution (p = .025 and .014 for the positive and the negative cluster, respectively). The late paired clusters showed a similar but polarity-reversed distribution at 326ms-475ms (p = .003 and .019 for the positive and the negative cluster, respectively.) These results are consistent with previous findings of EEG correlates of prediction for isochronous temporal sequences (Chen et al., 2010; Jacobsen & Schröger, 2003; Joutsiniemi et al., 1998). Note that, the latency of these responses, with respect to S2 onset, was 77ms (227ms – 150ms) and 176ms (326ms – 150ms) for the early and the late effects respectively. The remarkably early 77ms latency suggests an influence of anticipatory or predictive processing (Katz, 2014).

**Figure 2.**
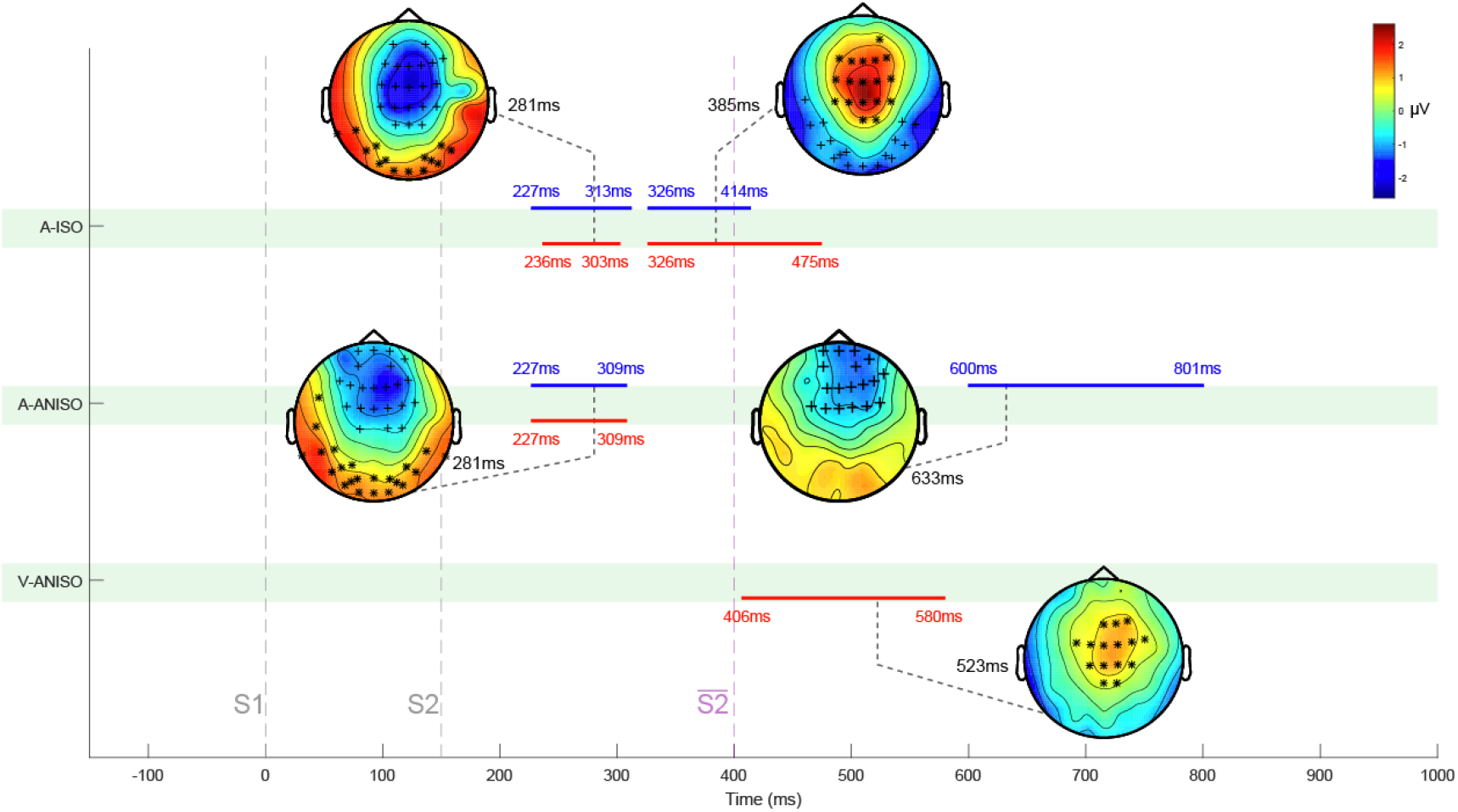
Differences in topography for mismatch response between standard and deviant 150ms durations. The temporal extent of statistically significant positive and negative clusters is depicted by red and blue lines respectively. The representative topographical distributions of each cluster are selected from representative time points within the time range of each clusters. Symbol * and + mark electrodes in positive and negative clusters, respectively, at representative time points. The two grey dashed lines indicate the timing of the first (S1, 0ms) and the second (S2, 150ms) stimuli in a trial. The purple dashed line indicates the expected S2 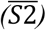 onset time (400ms) in deviant trials (i.e, when the 150ms duration is unexpected). A-ISO: Auditory isochronous sequence. A-ANISO: auditory anisochronous sequence. V-ANISO: visual anisochronous sequence.

We next examined whether EEG patterns across channels contained information about trial types, using MVPA. This approach generally shows improved sensitivity relative to univariate ERP analyses (Blankertz, Lemm, Treder, Haufe, & Müller, 2011; Stokes, Wolff, & Spaak, 2015; Wolff, Ding, Myers, & Stokes, 2015). Accordingly, as shown in Figure 3 (A-ISO), a linear SVM could classify standard and deviant EEG patterns with above chance performance, in two extended time windows. The later time window showed a longer range from 203ms to 805ms (p < .002) than the corresponding ERP analysis (326 - 475ms; A-ISO) suggesting that information about trial type (standard or deviant) was embedded across multiple electrode channels in ways not captured by univariate analyses.

**Figure 3.**
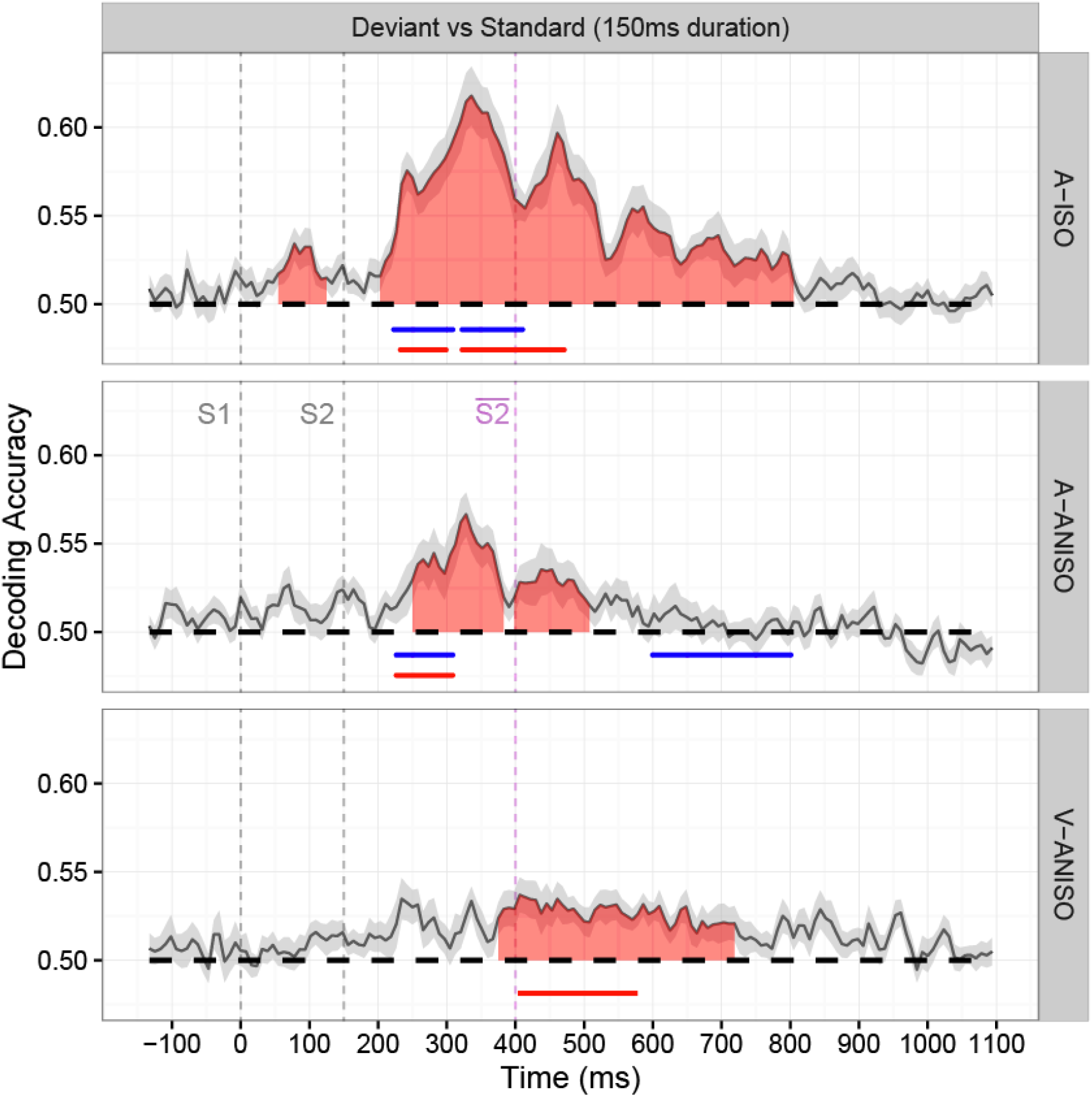
Average decoding accuracy across all participants for standard versus deviant trials of 150ms duration. Linear SVM classifiers were trained and tested to distinguish standard and deviant durations with data from the 64 electrode channels. Highlighted red areas show statistcally significant time windows by cluster-based permutation with 1D (time) data. The two grey dashed lines indicate the timing of the first (S1, 0ms) and the second (S2, 150ms) stimuli in a trial. The purple dashed line indicates the expected S2 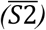 onset time (400ms) in deviant trials (i.e, when the 150ms duration is unexpected). Red and blue horizontal lines indicate the temporal extent of ERP mismatch, as shown in Figure 2. Shaded areas around the mean represents standard error of the mean decoding accuracy (SEM) across subjects. A-ISO: Auditory isochronous sequence. A-ANISO: auditory anisochronous sequence. V-ANISO: visual anisochronous sequence.

We also found a remarkably early significant window (p =.03) ranging from 55ms to 125ms (from S1 onset). Note that this time range is earlier than the appearance of the S2 that would mark the difference between standard and deviant trials (i.e., 150ms). Therefore, the early EEG pattern difference cannot be attributed to any violation of duration prediction, but was likely due to differences between short and long standard duration trials in the time-course of neural preparation for the predicted S2, facilitated by the rhythmic structure of isochronous sequences.

#### Prediction violation in anisochronous auditory sequences

Having replicated, and extended via MVPA analyses, previous results reported for rhythmic (isochronous) sequences, we examined physically identical short durations (150ms) when presented as standard or deviant in anisochronous stimulus sequences that, by design, were free from global rhythmic structure.

For the A-ANISO sequences, the ERP analysis (Figure 2, A-ANISO) revealed a pair of significant early clusters (ps < .002 for both positive and negative clusters) and a late negative cluster (600ms to 801ms, p < .002). These ERP differences between standard and deviant trials demonstrate that neural correlates of prediction can be found for duration itself, in isolation from rhythmic temporal information. Similar to the results from the A-ISO sequences, a significant early cluster pair showed that unexpected short durations elicited a central-to-frontal negative response at 227ms and 309ms. The early onset of these clusters suggests that prediction about duration influenced very early auditory processing, occurring only 77ms from S2 onset time (227ms - 150ms = 77ms). Differences between A-ISO and A-ANISO conditions appeared for later components. Unlike A-ISO sequences, ERP analysis of A-ANISO sequences did not show evidence for a second cluster (~ 300ms - 400ms), suggesting that this cluster in the A-ISO data reflects a unique contribution from rhythmic information to duration prediction. In combination with the results from presentation of isochronous sequences, these data indicate that both duration and rhythmic predictions modulated early auditory processing. Additionally, we found a very late (600 – 800ms) central-to-frontal negative cluster only in A-ANISO sequences, suggesting a unique contribution from duration, not found when the duration is embedded in a rhythmic sequence.

Next, linear SVMs were used to decode EEG patterns for A-ANISO sequences (MVPA analysis). The results show two time windows with above chance decoding accuracies. The early time window is consistent with the early ERP cluster and extends the time range (250ms to 383ms, p < .002, vs., 227 - 309ms for the ERP analysis). As with the A-ISO analysis, this suggests that MVPA is more sensitive to the neural response differences between standard and deviant trials than the ERP analysis. The later time window occurred at 398ms to 508ms (p < .011). We note that these windows are separated by only a brief period (15ms) of below statistically significant performance, suggesting that the window may in fact be continuous as in the A-ISO results. Notably, we did not find any early time window in which the MVPA performed above chance, by comparison to the ~55-125ms in the A-ISO condition, providing further evidence that this early window of decoding performance in the A-ISO condition relies on predictions based on rhythmic structure, not duration.

#### Prediction violation in anisochronous visual sequences

We next applied the same ERP and MVPA analyses as above to the V-ANISO condition, to examine the neurophysiological correlates of duration prediction for rhythm-free *visual* sequences. The ERP cluster-based analysis revealed that unexpected short durations (i.e., S2 at 150ms) elicited a significant mismatch cluster between ~406ms to ~580ms (p < .002), localized in central and frontal areas. As before, MVPA revealed a much longer significant time window, with above chance decoding accuracies from ~375ms to ~719ms (p < .002). Compared to auditory sequences, both ERP and MVPA analyses reveal a substantially longer latency for visual mismatch responses, consistent with differences in information processing speed between auditory and visual systems (A. J. King & Palmer, 1985; Regan, 1989).

### Generalization of duration prediction across modality

We examined whether the neural responses related to violations of duration prediction elicited by auditory and visual sequences contained shared information that may be associated with supramodal processing. We reasoned that if duration prediction is accomplished by a supramodal neural process, this could be revealed by the generalization of EEG pattern classification across modalities; specifically, a pattern classifier trained to decode auditory standard from deviant trials using EEG signals should also be able to decode visual standard from deviant trials. However, as shown above (Figures 2 and 3), there are inherent processing latency differences between auditory and visual modalities (auditory was always faster than visual) which may undermine simple tests of generalisation. Therefore, we used temporal generalisation analysis (TGA; J.-R. King & Dehaene, 2014). TGA assesses the generalisation of classification across time. If a classifier trained to differentiate two conditions (e.g. a standard vs. deviant duration) based on EEG signals at a specific time point (training time) is able to classify EEG patterns at a different time point (testing time) with an above chance accuracy, the EEG signals at the two time points are likely to contain shared information about the classification (J.-R. King & Dehaene, 2014). This method can be naturally extended to assess shared information not only across time but also across modality.

To examine whether shared information was embedded in neural patterns elicited by the visual and auditory short duration (150ms) deviants we applied TGA. At every sample point (training time) in the epoch range (-140ms to 1400ms), linear SVM classifiers were trained to classify neural patterns elicited by short duration (150ms) *auditory* standard and deviant anisochronous sequences (A-ANISO). These classifiers were then used to classify EEG signals at every time point (testing time) in *visual* anisochronous (V-ANISO) short duration standard and deviant trials. Repeating this training and testing procedure for all combinations of training and testing time-points creates a generalisation matrix (Figure 4d). Inspection of this matrix reveals a below-diagonal cluster having above chance classification accuracies (p =.021, Figure 4d). The existence of this cluster implies that information used to decode standard and deviant trials in short duration anisochronous auditory sequences can also be found in the neural response patterns elicited by visual sequences, though shifted later in time. Similar results were also found when we reversed the training and testing data sets (train on V-ANISO and test on A-ANISO). As expected, the generalisation matrix showed an above-diagonal cluster (p =.025, Figure 4b**Error! Reference source not found**.), indicating that the information used to decode neural responses elicited by standard and deviant short duration visual trials could successfully classify those in short duration audio trials, though at earlier time-points.

**Figure 4.**
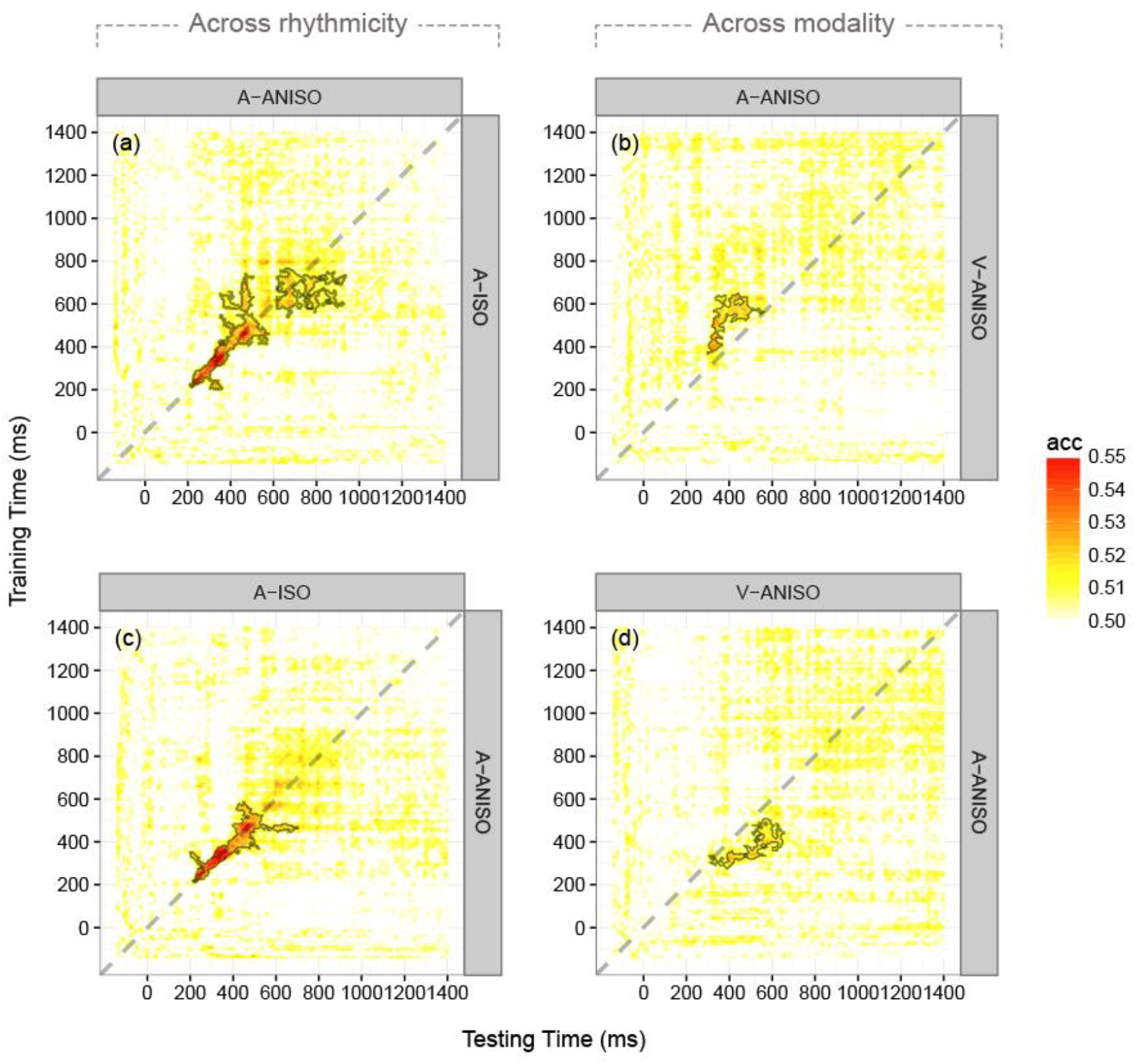
Temporal generalisation analysis (TGA) for classifying deviant vs standard 150ms durations. Classifiers were trained for each subject on EEG data at each training time-point and tested on data at each testing time-point. Panels on the left (a and c) show decoding accuracy for across-condition TGA comparing the isochronous (A-ISO) and anisochonous (A-ANISO) auditory conditions. Panels on the right (b and d) show decoding accuracy for across-modality TGA comparing the anisochonous auditory (A-ANISO) and visual (V-ANISO) conditions. Contour outlines indicate statistically significant clusters. Within each panel, clusters along the dominant diagonal (grey dashed line) show that EEG signals at nearby training and testing time-points share similar information. Clusters below the diagonal indicate that classification-relevant information at a training time-point is common with a later testing time while above diagonal clusters indicate that information at a training time-point is common with an earlier testing time.

To quantify the temporal offset in decoding performance between auditory and visual stimuli, we computed the average delays (average of the testing time minus the training time for each time point within a cluster) within the significant clusters. The TGA trained on audio and tested on visual showed mean latency difference of 131ms (t(399) = 42.52, p < 10^-149), while that trained on visual and tested on audio showed a mean difference of -142ms (t(387) = -46.07, p < 10^-158).

### Generalization of prediction across isochronous and anisochronous sequences

Finally, as when checking for commonalities in response across visual and auditory modalities, we used TGA to look for common periods of neural response to duration prediction violation between the two auditory conditions, one containing rhythmic information (A-ISO) and the other being arrhythmic (A-ANISO). We first examined whether information used to decode EEG patterns in the A-ANISO sequence could also decode EEG patterns in A-ISO sequences, even with the involvement of rhythmic-related processes. The results showed a significant cluster along the diagonal when trained on data from the A-ISO sequence and tested on data from the A-ANISO sequence. The decoders could classify EEG patterns (p < .01, Figure 4a) with above chance decoding accuracies. Note that the period of statistical significance along the diagonal was from 219ms to 523ms, suggesting a similar dynamic of EEG patterns shared between isochronous and anisochronous sequences within this period. The reverse TGA (trained on data from A-ISO and tested on A-ANISO) also showed a significant cluster with a similar generalisation pattern and time range (diagonal significant period from 219ms to 516ms, p < .01, Figure 4c), suggesting robust common neural activity in response to violations of expected duration for A-ISO and A-ANISO sequences during this period.

## Discussion

We sought to identify neural responses underlying duration prediction when presented in rhythmic or arrhythmic sequences, in auditory and visual modalities. We analysed EEG recorded during an extended temporal oddball paradigm using a combination of univariate (ERP cluster-based permutation analysis) and multivariate (multivariate pattern analysis, MVPA, and temporal generalization analysis, TGA) approaches. Our results revealed clear signatures of violation of duration prediction, both in the presence and absence of rhythmic structure. Regardless of whether a sequence did or did not contain rhythmic structure, presentation of an unexpected duration yielded a deviation in EEG responses similar to previous findings using only rhythmic sequences (Chen et al., 2010; Joutsiniemi et al., 1998; Näätänen et al., 1989; Näätänen, Paavilainen, Rinne, & Alho, 2007). The similarity in neural responses to violations of rhythmic and arrhythmic auditory duration predictions apparent when comparing the ERP and MVPA analyses across these conditions (Figure 2 and 3 A-ISO versus A-ANISO) was supported by the direct contrast of neural responses using temporal generalization analysis (TGA). Importantly, beyond these common neural signatures, we also identified distinct components of neural responses related to prediction of rhythm, and duration itself. Neural signatures of duration prediction specific to rhythmic sequences reflected very early anticipatory auditory processing (Figure 3 A-ISO), while violation of duration predictions in arrhythmic sequences was indicated by very late processes, concentrated over central to frontal brain regions (Figure 2 A-ANISO), and possibly related to prediction updating (Cohen, Wilmes, & van de Vijver, 2011; Mas-Herrero & Marco-Pallarés, 2014; Philiastides, Biele, Vavatzanidis, Kazzer, & Heekeren, 2010). These results are at least partially consistent with a previously suggested distinction between *duration-based* and *beat-based* timing (Grube, Cooper, Chinnery, & Griffiths, 2010; Grube, Lee, Griffiths, Barker, & Woodruff, 2010; Teki, Grube, & Griffiths, 2012; Teki, Grube, Kumar, & Griffiths, 2011). Overall, our results provide evidence for the existence of explicitly *duration* based processes for temporal prediction, independent of rhythmic structure.

It has long been debated whether duration information is processed in a modality specific or a modality-general manner (Buonomano & Karmarkar, 2002; Grondin, 2010; Ivry & Schlerf, 2008; N’Diaye, Ragot, Garnero, & Pouthas, 2004; Shih, Kuo, Yeh, Tzeng, & Hsieh, 2009; van Wassenhove, Buonomano, Shimojo, & Shams, 2008; Wearden et al., 2006). Using TGA, we found that classifiers trained to differentiate standard and deviant durations in auditory sequences could also decode standard and deviant durations in visual sequences (and vice-versa). Specifically, we showed similar, though temporally offset (vision delayed relative to audio), neural response patterns were elicited by unexpected short durations in auditory and visual modalities. These data suggest a shared supramodal component to duration prediction, operating across sensory modalities. Whether this means that duration processing itself is similarly shared, or that modality-specific duration processes act as input to the same supramodal predictive mechanism is an issue of interest for future study.

When assessing neural responses to duration deviants within a given modality, our data from both ERP and MVPA analyses showed longer response latency in response to violations of prediction for visual duration than for auditory (Figure 2 and 3, A-ISO and A-ANISO versus V-ANISO). TGA analysis permitted greater insight into these latency differences. When applying classifiers trained in one modality to data from the other, TGA showed average decoding latency differences of 131ms (when trained on auditory and tested on visual) and -142ms (when trained on visual and tested on auditory), with responses to prediction violations in vision always lagging those in audition (Figure 4b and 4d). These data show that neural responses encoding a supramodal distinction between standard and deviant intervals can be found at least 131ms earlier for auditory stimuli than for visual stimuli. Interestingly, basic neural responses associated with auditory signals can be found roughly 30ms to 50ms earlier than for visual signals (A. J. King & Palmer, 1985; Regan, 1989). This is notably shorter, by about ~100ms (longer for vision), than the across-modality latency revealed in our TGA analysis. One possible explanation for this difference is that the generation or comparison of predictions for visual durations may take longer than for auditory durations. However, in this study we did not attempt to equate the saliency and intensity level of the stimuli across the two modalities, and properties like stimulus intensity of have been shown to affect sensory information processing speed (Nissen, 1977; Töllner, Zehetleitner, Gramann, & Müller, 2011). Further research is needed to evaluate the impact of stimulus saliency and modality on interval prediction across modalities.

Neuroimaging studies have benefited in recent years from the development of multivariate analyses, like MVPA and TGA, to complement traditional univariate approaches. In our study, the two approaches provided generally consistent results. MVPA (and by extension TGA) generally displayed a greater sensitivity when distinguishing deviants from standards, than corresponding univariate approaches, leading to some novel insights. For example, for isochronous presentations, MVPA suggested the presence of a very early predictive component (starting as early as ~ 55ms after S1, which is almost 100ms prior to the predicted timing of the second event S2). Since S2 had not occurred at this point, this early component can best be interpreted as a prediction based on the rhythmic structure of the sequence. By comparison, the univariate ERP analysis showed no evidence for this rhythmic prediction. Additionally, temporal windows of statistical differences between the deviants and standards were generally longer for MVPA than for ERP. Under isochronous presentation, significant ERP differences extended from ~ 230 to ~ 470ms following S1, while for MVPA the window extends from ~200 to ~800ms. This increase in sensitivity is consistent with the idea that MVPA can take account of spatially distributed information across the whole array, while univariate ERP analyses do not.

Traditionally, comparison across conditions and modalities in EEG data is complicated by differences in sensory processing latencies associated with the different presented stimuli. As discussed above, this is particularly problematic for comparison across sensory modalities due to the well-established latency differences between visual and auditory processing streams. Comparisons using standard ERP or MVPA analyses must rely only on superficial correlations between time courses of neural responses evident in each individual ERP or MVPA analysis. By contrast, the use of TGA enabled the direct comparison of signatures of prediction violation in the isochronous (A-ISO) and anisochronous (A-ANISO) auditory sequences, by testing classifiers trained in one modality (across multiple time points) on the other (again across multiple time points). Our comparison of the patterns of neural activations associated with prediction violation across different sensory modalities was likewise facilitated by the use of TGA. While further research is needed to fully understand the similarities and differences of prediction and prediction violation across sensory modalities, it is important to note that the initial observations reporter here would have been impossible using traditional ERP or MVPA analysis methods.

Our results are broadly consistent with a predictive processing account of perception (Clark, 2012; Friston, 2005) which suggests that perceptual systems encode regularities in sensory inputs and use these regularities to make predictions about upcoming sensory signals. Ongoing comparison of bottom-up sensory inputs with top-down predictions produces prediction errors – the difference between the predicted and actual sensory data. The prediction error is then used to update internal models encoding the input regularities and predictions. Several studies have suggested that EEG components related to prediction violation, such as the MMN, might constitute empirical measures of prediction error in the framework of predictive processing (Garrido et al., 2008; Lieder, Daunizeau, Garrido, Friston, & Stephan, 2013; Lieder, Stephan, et al., 2013). By revealing the electrophysiological signatures of prediction violation for duration, that are independent of rhythmic structure and which generalize across modality, our study suggests a potential neural basis for predictive processing of duration perception. Furthermore, since time (in terms of duration) is a more abstract property than primary sensory features such as pitch or luminance, our data are consistent with the idea that predictive processing provides a general principle for perception across many dimensions of input modality and hierarchical organisation.

Overall, our findings demonstrate that humans can acquire predictions specifically related to duration, in the absence of rhythmic information, in both visual and auditory modalities. We further identified a set of neuronal signatures that reflect violations of duration prediction regardless of stimulus presentation modality, indicating that some aspects of duration prediction occur supramodally. Altogether, these findings suggest that duration as a property of sensory events can be encoded and used to generate predictions, and further, that temporal predictive mechanisms are shared across sensory modality, consistent with a predictive processing framework for perception.

## Acknowledgments

We thank David Schwartzman for advice and comments on the manuscript. Acer Y.-C. Chang was supported by a graduate teaching assistantship from the School of Engineering and Informatics at the University of Sussex, by a scholarship provided by Sackler Centre for Consciousness Science, and by the ministry of education Taiwan. Anil K. Seth and Warrick Roseboom are supported by the Dr.Mortimer and Theresa Sackler Foundation and the EU FET Proactive grant TIMESTORM: Mind and Time: Investigation of the Temporal Traits of Human-Machine Convergence.

